# Common framework for “virtual lesion” and state-dependent TMS: the facilitatory/suppressive range model of online TMS effects on behavior

**DOI:** 10.1101/187005

**Authors:** Juha Silvanto, Zaira Cattaneo

**Affiliations:** University of Westminster, Faculty of Science and Technology, Department of Psychology, 309 Regent Street, W1B 2HW, London, UK; Department of Psychology, University of Milano-Bicocca, 20126 Milan, Italy; Brain Connectivity Center, National Neurological Institute C. Mondino, 27100 Pavia, Italy

## Abstract

The behavioral effects of Transcranial Magnetic Stimulation (TMS) are often nonlinear; factors such as stimulation intensity and brain state can modulate the impact of TMS on observable behavior in qualitatively different manner. Here we propose a theoretical framework to account for these effects. In this model, there are distinct intensity ranges for facilitatory and suppressive effects of TMS: low intensities facilitate neural activity and behavior whereas high intensities induce suppression. The key feature of the model is that these ranges are shifted by changes in neural excitability: consequently, a TMS intensity, which normally induces suppression, can have a facilitatory effect if the stimulated neurons are being inhibited. For example, adaptation reduces excitability of adapted neurons; the outcome is that TMS intensities which inhibit non-adapted neurons induce a facilitation on adapted neural representations, leading to reversal of adaptation effects. In conventional virtual lesion paradigms, similar effects occur because neurons not tuned to the target stimulus are inhibited. The resulting reduction in excitability can turn high intensity inhibitory TMS to low intensity facilitatory TMS for these neurons (whereas neurons tuned to the target stimulus are inhibited), leading to a reduction in signal-to-noise ratio. Thus differential excitability levels of neural populations contributing to behavior, combined with nonlinear neural effects, can explain how TMS modulates behavior.

## 1. Introduction

The era of using transcranial magnetic stimulation (TMS) to study perceptual and cognitive functions began with the classic work of Amassian et al (1989) who showed that applying TMS over the visual cortex impaired participants’ ability to report briefly presented letters. Furthermore, in what was the earliest demonstration of the focality in TMS studies of visual perception, Amassian et al (1989) demonstrated that when the coil was moved slightly laterally, most disrupted were letters that fell into the contralateral hemifield. This work was important for demonstrating how TMS can be used to study perceptual functions with an excellent temporal resolution and with a reasonable spatial specificity. But it was important also in another manner: it arguably defined our way of thinking about TMS as a *disruptive* tool – for inducing “virtual lesions”. In other words, it paved the way for conceptualizing TMS as a “lesion” technique, with impairment as its default outcome. In this context, facilitations would not be expected – and when they have occurred, they have often been referred to as “paradoxical” (e.g. Fecteau et al, 2006; Theoret et al, 2003).

More recent studies have shown the situation to be much more complex and the view of TMS as a disruptive technique does not hold in a wide range of circumstances. Factors such as stimulation intensity, task difficulty and cognitive state can fundamentally change the nature of behavioral TMS effects (see e.g. Sandrini et al, 2011; de Graaf et al, 2014; Romei et al, 2016 for reviews). In this review we will briefly discuss how factors such as stimulation intensity and brain state can qualitatively modulate the direction of TMS effects. We will then provide a conceptualization of these effects which includes both standard and state-dependent TMS paradigms. Our discussion is focused on online TMS paradigms in which single-pulses or brief pulse trains are applied concurrently with the behavioral task; thus “offline” paradigms (such as theta burst or 1 Hz rTMS) which induce longer-lasting aftereffects are beyond the present discussion.

## 2. The importance of stimulation intensity in determining behavioral effects of TMS

In the study of Amassian et al (1989), participants were presented with trigrams of randomly chosen letters, briefly presented at fixation. Single-pulses of TMS were applied over the calcarine cortex (2 cm above the inion at midline) at time windows ranging from 0 and 200 ms after stimulus offset. When TMS was applied 80-120 ms after stimulus onset, participants’ ability to detect the letters was impaired. When interpreting these results, it is interesting to consider the parameters with which the effects were obtained: single pulses of TMS were applied over the occipital cortex at a high intensity, 90-100% of the maximum output of a Cadwell stimulator with a circular coil (2.2 T) (but note that the link to phosphene thresholds is unclear due to lack of phosphene induction in that study). Behaviorally, baseline performance was close to ceiling. The use of such parameters makes sense – when aiming to modulate behavior with TMS, it sounds reasonable to maximise the physical level of an intervention aimed at modulating behavior, while having a reasonable high level of baseline performance. Subsequent literature on visual masking by TMS has shown that intensity indeed matters: the intensity level needed to impair the visual perception of external stimuli is relatively high, for example in comparison to the intensity needed to induce a phosphene threshold (PT) (e.g. Kammer, 2005; see Kammer, 2007, de Graaf et al, 2014, for review). Thus in general, the use of TMS in cognitive neuroscience has involved the use of relatively high threshold intensities (relative to PTs).

Common to the studies described above is the combination of high TMS intensity (relative to phosphene threshold) and high level of baseline performance. Interestingly, with lower intensities TMS has been shown to *facilitate* performance (e.g., Abrahamyan et al, 2011, 2015; Schwarzkopf et al, 2011). In a study by Schwarzkopf et al (2011), participants were asked to perform a motion direction discrimination task which they were thresholded to perform at either low (60%) or high (85%) baseline level. Concurrently with stimulus onset, a TMS pulse train (three pulses at 10 Hz) was applied over V5/MT at different intensities. The key finding was that, whereas high intensity TMS impaired performance when baseline task performance was high, low intensity TMS facilitated motion discrimination when baseline level of performance was low. Facilitations by subthreshold TMS have also been found by a series of studies by Abrahamyan et al (2011, 2015). In their 2011 study, single pulses of subthreshold TMS were applied over the early visual cortex (within the time window in which suprathreshold TMS impairs detection) while the contrast of test stimuli was varied using an adaptive staircase procedure. The results showed a decrease in contrast sensitivity.

## 3. Nonlinear TMS effects as a function of brain state

Another example of nonlinear TMS phenomena comes from manipulations of brain states prior to TMS application. State-dependent paradigms differ from conventional “virtual” lesions in that, in the former, preconditioning is used to modify the activity state of neuronal populations before TMS is applied (see e.g. Siebner et al, 2004; Silvanto et al, 2008; Romei et al, 2016 for reviews). One such effect has been observed in studies in which participants are adapted to a specific visual attribute (such as color or direction of motion) prior to the application of brief pulse trains of TMS. In adaptation, prolonged exposure to a sensory stimulus (such as colour, shape or motion) can lead to perceptual aftereffects in which perception is biased away from the exposed stimulus. For example, viewing a motion stimulus moving leftward can induce subsequently presented stationary stimulus to appear moving rightward. The central finding in these studies is that TMS abolishes or even reverses the impact of adaptation (e.g. Stewart et al, 1999; Theoret et al, 2002; Campana et al, 2011, 2013, Silvanto et al, 2007, Guzman-Lopez et al, 2011). For example, in a study by Silvanto et al (2007), participants were adapted to a uniform red colour for 30 seconds, after which phosphenes were induced from the early visual cortex. Counterintuitively, the phosphenes took on the colour of the adapting stimulus. Similar results were obtained in a further experiment in which participants were adapted to a conjunction of orientation and colour; after adaptation, TMS reversed the adaptation effect such that detection of items identical to the adapter was superior to detection of items which differed from the adapter in both colour and orientation. Subsequent studies have found state-dependent effects in adaptation paradigms in a range of domains, including motion direction discrimination (Cattaneo & Silvanto, 2008; see also Campana et al, 2002, 2006 for earliest TMS-priming studies), number processing (Cohen Kadosh et al, 2010; Renzi et al., 2011), action observation (Cattaneo, Sandrini & Schwarzbach, 2010; Jacquet & Avenanti, 2015) and emotion perception (Mazzoni et al, 2017). In addition to adaptation, preconditioning by priming has been used to induce similar effects (Cattaneo et al, 2008; Cattaneo, Silvanto, et al. 2009; Cattaneo L, 2010). Puzzlingly, such facilitations have been found with suprathreshold intensities which in the absence of adaptation impair behavior.

## 4. Why an explanation in terms of “noise” cannot account for TMS effects

While TMS effects are generally described in terms of “interfering” or “disrupting” behavior, there have been attempts to explain its effects on behavior more mechanistically, in terms of inducing noise. The idea is that TMS *indiscriminately* activates neurons in a targeted region and in this manner adds noise to neural processing. This noise reduces the signal-to-noise ratio of the cognitive task under investigation and thus impairs performance (Walsh & Pascual-Leone, 2003; Miniussi et al, 2013). In this view, TMS intensity is equated to the amount of noise added to neural processing.

The noise model can explain facilitations by TMS described above. An important aspect of “noise” is that it is not always detrimental to behavior – this depends on the amount of noise and initial signal strength. In systems with measurement threshold, the addition of low levels noise can in fact push weak subthreshold signals beyond the threshold, improving information transfer. This is known as *stochastic resonance* (Stocks et al, 2000). The key issue is level of noise - when noise level is too high, the signal is weakened too much. The studies discussed above provide evidence that there are such stochastic resonance effects in TMS. This can explain the findings discussed above (Schwarzkopf et al, 2011; Abrahamayan et al, 2011) where low intensity TMS had a facilitatory effect on motion discrimination when the baseline performance level was relatively low. In this view, low intensity TMS would be adding low levels of noise to a weak signal that is near threshold; this would push activation above threshold. When TMS intensity is increased, the level of noise drowns out the signal.

There are however critical limitations of the noise models. Firstly, they cannot explain various behavioral nonlinear TMS effects, for example those observed with adaptation. According to the “noise” model, TMS would indiscriminately stimulate all neurons regardless of brain state, adding random noise to both adapted and nonadapted neurons. It is difficult to see how this could lead to the activity of adapted populations to dominate neural read-outs such that behavioral adaptation effects are reversed; high levels of noise would lead to saturation of all neuronal populations. One might argue that the amount of “noise” being added by TMS may vary between neuronal populations, but this is inconsistent with the concept noise being statistically independent of signal-induced activity (and would still struggle to explain behavioral reversals). It is in fact difficult to explain state-dependent TMS effects without postulating *a differential TMS effect* on neurons in different states, such as preferential activation of adapted neural populations, as has been proposed previously (e.g. Silvanto et al, 2008). But this raises the question of how to explain the wide range of TMS effects within a common framework.

A further issue relates to the neural plausibility of conceptualizing the impact of TMS merely as noise. An issue raised by various groups is that differential effects of TMS may reflect the saturation of activity among neurons that are already responding to other inputs (such as a sensory signal). For example, less active neurons may be more strongly excited by TMS than more active neurons (e.g. Silvanto & Muggleton, 2008, Ruzzoli et al, 2011). But if this is the case, then TMS cannot be thought in terms of adding random noise, as “noise” by definition should be statistically independent of signal-induced activity (point made previously by Ruzzoli et al, 2011). Thus view of TMS simply adding noise therefore appears to be too simplistic.

Below we propose a framework which differs from the “noise” models in that the impact of TMS is not indiscriminate; rather, the effect of a TMS pulse can differ greatly between neuronal populations, as a function of their neural excitability levels. Indeed, differences in excitability of neurons contributing to behavior play a key role in determining the behavioral outcome of TMS.

## 5. Explaining TMS effects in terms of facilitative and suppressive ranges of stimulation

The model is rooted in evidence from studies on the visual system. However, before discussing its details, it is important to address the issue of how neural and behavioral effects of TMS can be linked. It might be argued that behavioral effects of TMS must be explained at behavioral level and care must be taken when linking neural activity to perception, with statements such as “TMS was used to suppressed neural activity to impair behavior” problematic due to mixing different levels of explanation. However, we would argue that it is not inherently false to make this link, *if sufficient information exists on the link between neural activity and behavior.* Such links often exist for low-level visual functions where developing models of neural readout underlying behavior are more straightforward than for higher level cognitive functions. For example, visual motion direction discrimination can be explained in terms of firing rates of direction-selective neural populations of different tunings, with computations such as vector averaging and winner-takes-all determining the perceptual report (cf. Nichols & Newsome, 2002). The key point is that there is a close link between neural activity level and behavior – allowing this link to be used to explain behavioral effects of TMS. Furthermore, because sufficiently is known regarding the neural effect of TMS in the visual cortex, the link between neural activity, its modulation by TMS, and behavior can be made.

To provide an example of this issue, let us consider the following argument in a hypothetical TMS study: “We hypothesised that *suppression* of V5/MT *impairs* motion perception”. Is one allowed to make the link between the putative TMS-induced suppression and a behavioral impairment induced by TMS? In our view this is justifiable, because the relationship between neural activity and TMS on one hand, and neural activity and behavior on the other, are empirically supported. Specifically, there is evidence that 1 Hz rTMS suppresses neural activity and cortical excitability – this has been shown for example using phosphene thresholds (e.g., Boroojerdi et al., 2000). Thus the argument that V5/MT is suppressed by TMS is empirically supported. The second step involves the link between neuronal activity in V5/MT and motion perception, i.e. how the read-out of firing rate of V5/MT neurons underlies motion perception (Nichols & Newsome, 2002). These two pieces of evidence allow the bridge between suppression of V5/MT activity and impairment in motion perception to be made. With respect to higher-level functions, this level of explanation might be more problematic, if no clear model exists of how the function arises, and the neural mechanisms underlying them have not been comprehensively mapped. For example, with respect to short-term memory, there is an ongoing debate on whether maintenance occurs via firing rate or synaptic mechanisms (e.g. Stokes, 2015); thus linking neural activity to behavior would be problematic. By using the above logic – linking known neural effects to behavior, we construct a new model to explain behavioral effects of TMS. While motivated by adaptation effects, this model generalizes to behavioral effects in other paradigms, and importantly, generates new testable hypotheses and is therefore falsifiable. The model key aspects are detailed below.

### Low intensity TMS facilitates early neural firing, higher intensity TMS suppresses it; the former effect is linked to facilitation of perception, the latter to impairment

We argue that the existence of facilitatory/inhibitory ranges for behavioral effects of TMS as a function of stimulation intensity is best accounted for by a similar nonlinearity in the effect of TMS on neural activity, likely to reflect inhibitory and excitatory networks having different thresholds. Moliazde et al (2003) investigated the impact of single pulse of TMS applied over the visual cortex by recording single-cell activity from cat’s visual cortex. The key finding was that weak stimulation (<50% of TMS intensity) caused an early facilitation of spontaneous and visually-induced activity up to 200 ms after TMS pulse, followed by a late inhibition. In turn, higher TMS intensities increasingly evoked an early suppression of activity for 100–200 ms which cancelled the early part of facilitation evoked with lower stimulus strengths. This was followed by a delayed, prolonged and increased facilitation (up to 500 ms) and terminated in inhibition.

While the pattern of neuronal activity induced by TMS appears complex, there are strong reasons to link the early component of the neural effect (i.e. facilitation by low intensity and suppression by high intensity stimulation, respectively) to behavioral findings. Specifically, the early neural suppression with high intensities can be linked to disruptive behavioral effects of TMS, as suprathreshold (relative to phosphene threshold) intensities are needed to induce these, as discussed above. The early neural suppression underlying behavioral effects of (high-intensity) TMS is also consistent with the timing of TMS-induced masking of visual information. Such masking effects (induced by applying TMS over V1/V2) typically occur when TMS is applied at time windows of 80-120 ms after target onset but not at later time windows (e.g., de Graaf et al., 2014; Kammer, 2007), indicating that an early neural effect must underlie them. Thus both the timing and intensity-dependency of TMS-induced visual masking match with the early suppression of neural activity reported by Moliazde et al (2003). In contrast, as discussed above, subthreshold TMS intensities applied within the TMS-masking time window can have a facilitatory impact on detection, consistent with the early facilitation of neural activity reported by Moliazde et al (2003). Overall, the available evidence suggests that stimulation of visual regions below phosphene threshold level (i.e. low TMS intensity) facilitates both (early) neural activity and visual detection, whereas and above threshold (i.e. high intensity) stimulation suppresses (early) neural activity and impairs behavior.

One might raise the question of whether making a link from neural facilitations and inhibitions to behavioral effects is appropriate? At least for low-level perceptual functions (such as motion direction discrimination), neural activation levels contribute rather directly to behavior, with firing rates of neural populations summed or averaged as a function of their tuning (Nichols & Newsome, 2002). Given the good correspondence between the neural and behavioral effects in terms of their direction (facilitation or inhibition) as a function of intensity, as well as their early timing, we argue that linking the two is justified. Furthermore, one could argue that a realistic model of behavioral effects needs to take such neural nonlinearities into account.

### TMS impairs detection because excitability changes dissociate facilitatory/inhibitory ranges of neurons tuned to the target stimulus and those not tuned to target

How do these distinct facilitatory and inhibitory ranges explain behavioral effects of TMS in conventional “virtual lesion” paradigms? This is shown in Figure 1. Again, an example can be made of motion perception. Let us consider an experiment where the participant is required to discriminate the direction of a motion stimulus. Motion-sensitive cortical regions contain distinct neuronal populations tuned to different motion directions; when a given motion direction is presented, neurons tuned to the stimulus are strongly activated, whereas the activity of neurons not tuned to the stimulus are inhibited (in top panel of Figure 1, the neurons which are selective for the presented stimulus are referred to as *“neurons tuned to target”*, and those not selective for it are referred to as “neurons not tuned to target”). This activation difference (i.e. signal-to-noise ratio) underlies behavioral motion discrimination (see blue bars in the lower panels in Figure 1). If TMS is applied at a sufficiently high intensity, as shown in Panel B (i.e. within the inhibitory range), neurons tuned to the target stimulus are inhibited. Importantly, the due to their differential excitability state, the effect of TMS on neurons not tuned to the target stimulus is different, as these are being inhibited and are thus less excitable (e.g. Rust et al., 2006). This reduction of excitability is akin to reduction in TMS intensity – as the same level of stimulation has a weaker effect on neural activity. Thus the facilitatory and inhibitory ranges of TMS in Figure 1 are shifted towards the right for these neurons – higher intensities are needed to obtain the same neural effect. *The consequence of this shift is that, with the TMS intensity shown in Panel B, neurons not tuned to the target are now in the facilitatory range (whereas neurons tuned to the target are in the inhibitory range).* And as the activation level difference between these neuronal populations determines the signal strength (see bars with TMS coils above them in Panel B), what follows is a reduction in signal-to-noise ratio and the observer’s sensitivity to the stimulus. Thus differential excitability of neurons contributing to behavior is critical.

**Figure 1.**
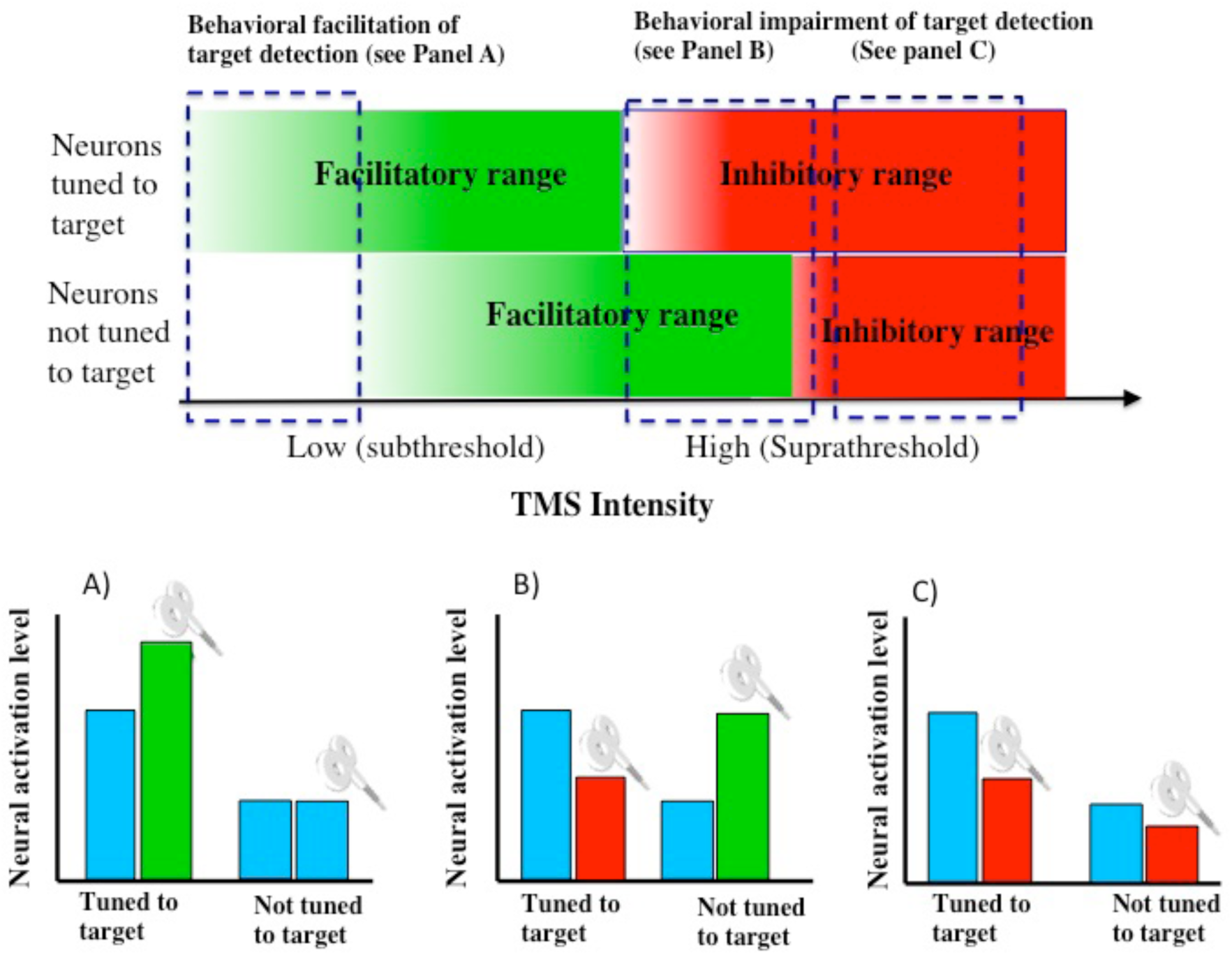
A model of TMS effects in a standard detection paradigm when TMS is applied concurrently with the target stimulus. The top panel indicates the facilitatory and inhibitory ranges for neuronal populations which are either tuned to or not tuned to the target stimulus. Panels at the bottom indicate the relative activation levels between these neuronal populations which underlies stimulus detection. When presented with a target stimulus, neurons driven by that stimulus are strongly active, whereas those not tuned to it are inhibited and thus less excitable. This reduction in excitability shifts the facilitatory/inhibitory range for non-tuned neurons to the right – as a higher TMS intensity is needed to activate them. At very low intensities, a facilitation in detection is expected, as neurons tuned to target are facilitated but intensity is too low to activate the inhibited neurons (see also Panel A at the bottom). Impairment of behavior occurs when neurons tuned to target stimulus are in the inhibitory range – the nature of the impairment depends on whether the inhibited neurons are in the facilitatory or inhibitory range (see Panels B and C).

In summary, the outcome of the reduced excitability of neurons not tuned to the target stimulus is a shift in the facilitatory/inhibitory range of TMS effects for these neurons - higher TMS intensity is needed to obtain the same neural effect. This gives to a rise situation in which *the same TMS intensity (shown in Panel B), facilitates these neurons while suppresses neurons tuned to the target.* This reduces or even reverses the activation difference between these neural populations which underlies perceptual sensitivity.

In contrast, At very low TMS intensities, a facilitation in stimulus detection can occur because neurons tuned to target stimulus are facilitated (i.e. they are in the low intensity facilitatory range) but the intensity is too low to activate neurons not tuned to the target stimulus, given the excitability reduction of the latter (see Panel A in Figure 1). This results in an increase in the activation level difference of the two neuronal populations (see the bars marked with a TMS coil in panel A) and thus an increase in signal-to-noise ratio. However, when both populations of neurons are in facilitatory range (i.e. both fall within the green region depicted in Figure 1), no change in behavior is predicted – as the relative activity level of the two neuronal populations would not drastically change.

### As adapted and non-adapated neurons have different profiles for facilitation and impairment, at certain TMS intensities former are facilitated and latter impaired: this reverses the impact of adaptation

State-dependent paradigms differ from conventional “virtual” lesions in that, in the former, preconditioning is used to modify the activity state of neuronal populations before TMS is applied. One such approach is the use of adaptation to preconditioning specific neural populations before application of TMS. When explaining TMS-adaptation effects, the key aspect to consider is the impact of adaptation on neural activity and susceptibility to external input. Adaptation induces a change in the neuron’s operating range (such as for contrast gain or sensitivity), with a stronger stimulus needed to induce a certain level of neural firing (Kohn & Movshon, 2003, (e.g., Kohn, 2007; Solomon & Kohn, 2014).). With respect to TMS, from this it follows that a higher TMS intensity is needed to induce firing in adapted neurons. Thus, the effect of adaptation for TMS-evoked responses is akin to turning down the TMS intensity - higher intensity is required to drive the neurons.

The counterintuitive effects of TMS after state-dependent manipulations such as adaptation are explained in Figure 2. “Adapted neurons” here refer to neurons which are strongly activated by the adapting stimulus and thus their excitability has been reduced, as discussed above (for example, after viewing rightward motion for a prolonged period, neurons tuned to this motion direction would be “adapted” whereas neurons tuned to other motion directions would be non-adapted). As discussed above, facilitatory and inhibitory effects of TMS operate at distinct intensity levels. With relatively low intensities, facilitations are obtained; with higher intensities, TMS has a suppressive effect. The key issue is that adaptation shifts these ranges, with the profiles for adapted and non-adapted neurons becoming diverged. This occurs because adaptation reduces neural excitability – and consequently, higher TMS intensity is needed to obtain the same neural effect as without adaptation (hence rightward shift in the top panel of Figure 2). Therefore, when TMS is applied at an intensity which normally impairs behavior, for adapted neurons this intensity now falls within the facilitatory range, due to the lower excitability of these neurons (see Panel B). In contrast, the non-adapted neurons are still in the inhibitory range. The behavioral outcome is the reversal or abolishment of adaptation effects, a shown in panel B. Specifically, whereas without TMS the activity level of nonadpated neurons is higher than that of adapted neurons (a difference which underlies adaptation aftereffects), inhibition of former and facilitation of latter reverses this pattern.

**Figure 2.**
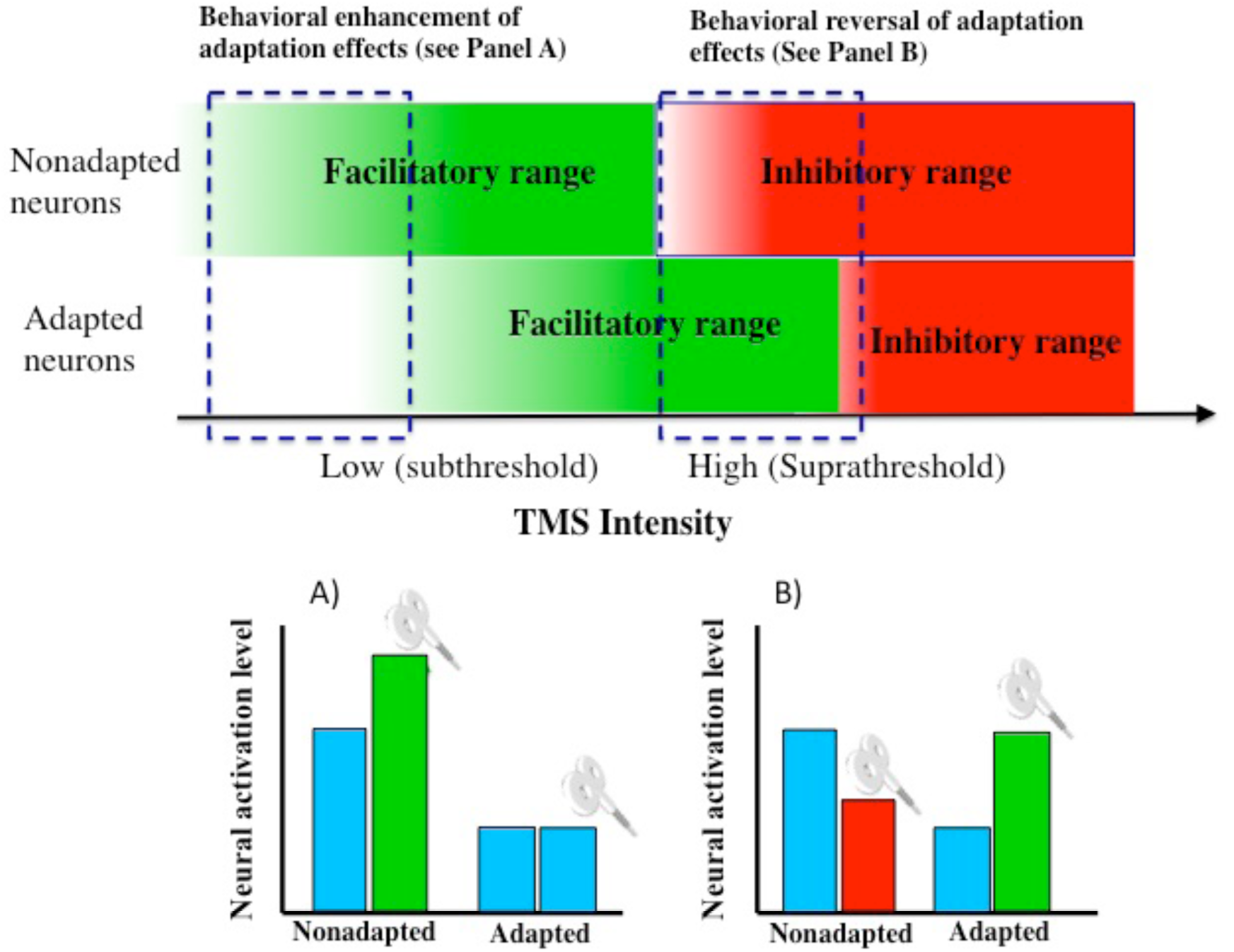
Explaining TMS-adaptation effects in terms of facilitatory and inhibitory ranges of TMS. Adaptation shifts the facilitatory and inhibitory ranges with respect of TMS intensity: following adaptation, when TMS is applied at an intensity at which non-adapted neurons are already in the inhibitory range, adapted neurons are still in the facilitatory range. Adaptation effects are reversed within this window (see panel B). In contrast, when TMS is applied at a lower intensity (see Panel A), an increase in the adaptation effect is predicted. This would happen because the low intensity TMS would enhance the activity of nonadapted neurons, whereas the adapted neurons (due to their loss of excitability) would not be driven by the TMS pulse.

An important point about this model is that the reversal/abolishment of adaptation is intensity-dependent. If TMS intensity is sufficiently low (see Panel A), one would observe an *enhancement* of adaptation effects. This would happen because the low intensity TMS would enhance the activity of nonadapted neurons, whereas the adapted neurons (due to their loss of excitability) would not be driven by the TMS pulse. An outcome is an increase in activation imbalance in favour of the nonadapted neurons. Thus if one were to parametrically vary TMS intensity, one would predict an enhancement of *adaptation* effects at the lowest intensities, a reversal of adaptation at intermediate intensities (i.e. with those generally used in TMS studies), and no effect on adaptation with highest intensities (as at highest intensities, both adapted and nonadapted neurons are suppressed and therefore their relative activity levels are not changed)

### Bringing together state-dependent and conventional TMS effects into a common framework

Figure 3 brings together the effects in conventional and state-dependent paradigms. The common theme is *neural excitability* and the effect of its modulation on facilitatory/inhibitory ranges of TMS Fundamentally, in all online TMS paradigms, the key issue in determining is the susceptibility of the neuronal populations to be activated by TMS. In state-dependent paradigms, manipulations such as adaptation and priming modulate this susceptibility. In conventional “virtual lesion” paradigms, excitability changes are induced by incoming information depending on the selectivity for the target stimulus: as discussed above, neurons not tuned to a target stimulus may be inhibited, leading to a change in excitability. Behavioral outcome is thus determined by the excitability of the various neuronal populations contributing to neural readout.

**Figure 3.**
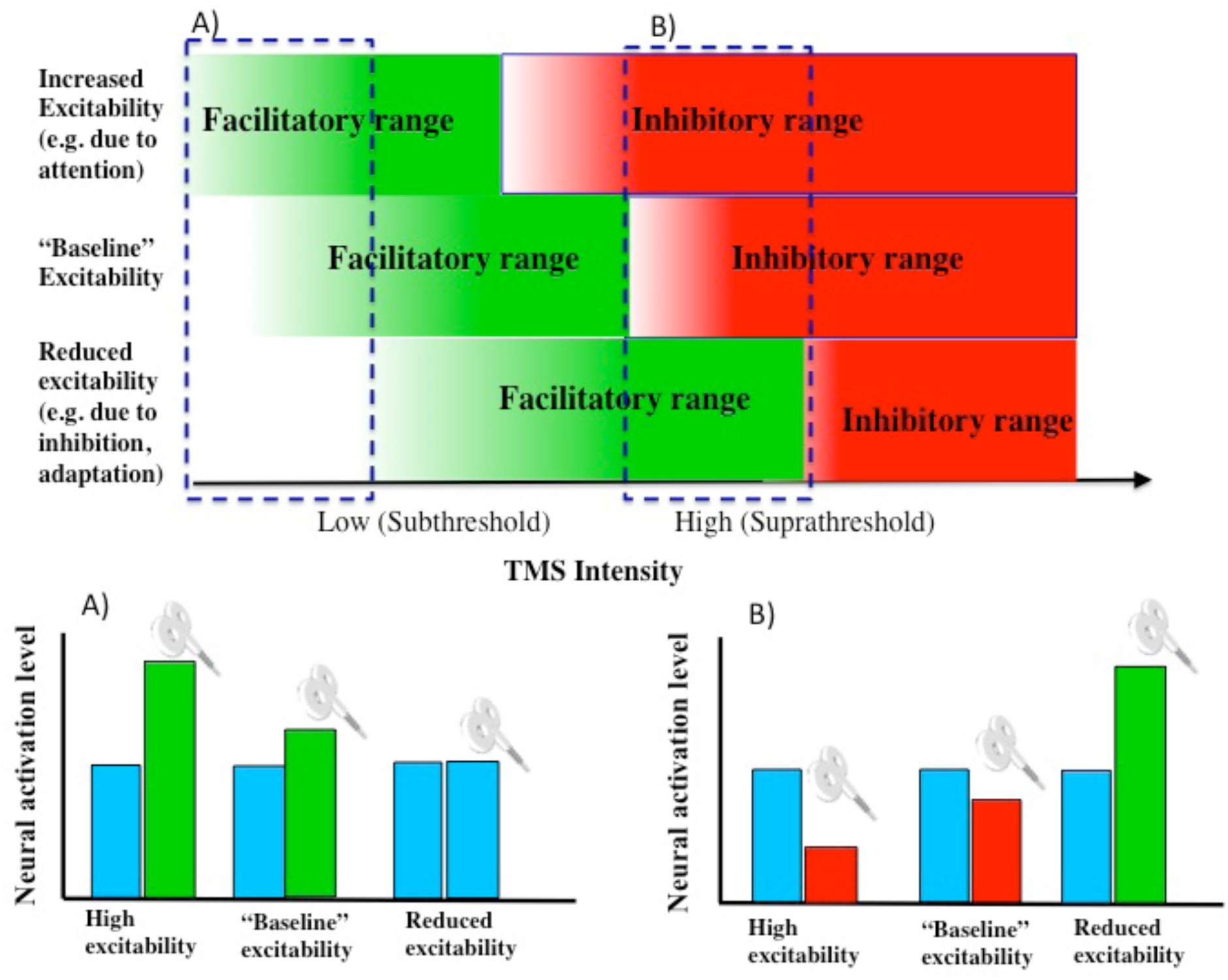
Changes in neural excitability shift the facilitatory/inhibitory range of TMS effects. TMS effects thus vary as a function of the excitability state of the neuronal populations contributing to behavior. A consequence is that a given set of TMS parameters can induce different behavioral effects depending on neural excitability at the time of stimulation. Note that in the bottom panel, for the purpose of demonstrating the impact of TMS, neuronal activation level in the absence of TMS is equal in the three levels of excitability. (This is not implausible as neural activation level and excitability are dissociable; e.g. Matthews, 1999.)

## 6. Testing the model: predictions

A key feature of any useful model is that it gives rise to predictions which can be empirically tested. These predictions relate to the nature of effects found at different TMS intensities. Testing of these predictions requires a systematic assessment of TMS effect from low to high intensities.

1) In conventional TMS paradigms, the model predicts that at the lowest subthreshold intensities, the neurons driven by a target stimulus are already in the facilitatory range whereas neurons not tuned to it are below the range where they can be facilitated (as they are inhibited – this shifts the facilitatory/inhibitory ranges rightwards). In this circumstance what we would see is a preferential activation of the “active” neurons - *the prediction is that target detection is facilitated.*

2) At intermediate subthreshold intensities, when both populations of neurons are in facilitatory range, *no change in behavior is predicted* – as the relative activity level of the two neuronal populations would not be greatly affected, due to both being in facilitatory range.

3) Impairment of target detection is obtained when neurons tuned to the target are in the inhibitory range. Interestingly, there is a stimulation window in which the neurons not tuned to the target are still in facilitatory range (see Panel B in Figure 1), whereas at highest TMS intensities, both neuronal populations are in the inhibitory range (see Panel C in Figure 1). We predict that these two intensities produce qualitatively different behavioral effects. Whereas in the former case, detection will biased towards the features encoded by items not tuned to the target (as these are still in the facilitatory window), in the latter case there will be a general of both neuronal populations. For this intensity, we would expect strongest behavioral effect in a task assessing the presence vs absence of motion, with TMS biasing responses towards the latter.

4) With respect to adaptation, reducing or increasing TMS intensity can lead to qualitative changes in adaptation effects. Reversal or abolishment of adaptation effects is only observed within the intensity range shown in Figure 2, i.e. when adapted neurons and non-adapted neurons are in facilitatory and inhibitory ranges, respectively. Thus if one were to parametrically vary TMS intensity, one would predict an enhancement of *adaptation* effects at the lowest **subthreshold** intensities, a reversal of adaptation at intermediate intensities (i.e. with those generally used in TMS studies), and no effect on adaptation with highest suprathreshold intensities (as at highest intensities, both adapted and nonadapted neurons are suppressed and therefore their relative activity levels are not changed).

## 7. Initial evidence for the model

The proposed model can explain not only the enhancements and impairments of perception observed in conventional TMS paradigms, but also the counterintuitive findings obtained with state-dependent manipulations, such as the reversal of adaptation and priming. To examine this issue further, we recently investigated (Silvanto, Bona, Cattaneo, in press) how state-dependent TMS effects depend on TMS intensity. We used a TMS-priming paradigm (see e.g. Cattaneo et al, 2008b), in which a visual prime (colour grating) was followed by a target stimulus which could be either congruent or incongruent with the prime. When TMS was applied in the time window in which V1/V2 TMS has been most efficient in modulating visual perception, TMS facilitated the detection of incongruent stimuli while not significantly affecting other stimulus types – a pattern of result generally found in TMS-priming studies (see e.g. Cattaneo et al, 2008, 2010). The key finding was that this effect was obtained only when TMS was applied at suprathreshold level (at 120% of phosphene threshold); subthreshold TMS (90% of PT) induced no effect.

In contrast, as discussed above, in the absence of state manipulations such as priming it is *subthrehsold* TMS rather than suprahtreshold TMS which facilitates visual detection (e.g. Abrahamyan et al, 2011). Thus when priming was used, *higher* stimulation intensity is needed to induce the same perceptual effect on stimuli incongruent with the prime. This is likely to reflect reduced susceptibility to TMS of neurons incongruent with the prime, consistent with the view that priming, by reducing neural excitability to incongruent targets, shifted the facilitatory/inhibitory range of TMS effects.

## 8. Concluding remarks

The proposed framework departs from the conventional “noise” view in arguing that, rather than indiscriminately adding random neural noise, the impact of TMS is strongly dependent on neural activation state. Specifically, the key to explaining TMS effects is to consider the different excitability levels of neural populations contributing to behavior (in combination with nonlinearities in neural effects of TMS). This is the case because neural excitability determines whether a given stimulation intensity falls into facilitatory or inhibitory range. Furthermore, it is important to stress that this model can be applied to behavioral TMS studies generally (rather than merely those which use preconditioning approaches); this is because excitability differences between neurons of different tunings arise whenever an external stimulus is being processed. While requiring empirical validation, a major potential strength of this model is its ability to offer a comprehensive framework for all online TMS studies.

## Acknowledgments

Juha Silvanto is supported by the ERC (336152).

## References

Abrahamyan, A., Clifford, C. W., Arabzadeh, E., & Harris, J. A. (2011). Improving visual sensitivity with subthreshold transcranial magnetic stimulation. The Journal of Neuroscience, 31(9), 3290–3294.

Abrahamyan, A., Clifford, C. W., Arabzadeh, E., & Harris, J. A. (2015). Low Intensity TMS Enhances Perception of Visual Stimuli. Brain Stimulation, 8(6), 1175–1182.

Amassian, V. E., Cracco, R. Q., Maccabee, P. J., Cracco, J. B., Rudell, A., & Eberle, L. (1989). Suppression of visual perception by magnetic coil stimulation of human occipital cortex. Electroencephalography and Clinical Neurophysiology/Evoked Potentials Section, 74(6), 458–462.

Boroojerdi, B., Prager, A., Muellbacher, W., & Cohen, L.G. (2000). Reduction of human visual cortex excitability using 1-Hz transcranial magnetic stimulation. Neurology, 54, 1529–1531.

Campana, G., Cowey, A., Walsh, V. (2002). Priming of motion direction and area V5/MT: a test of perceptual memory. Cereb.Cortex 12(6),663–669.

Campana, G., Cowey, A., Walsh, V. (2006). Visual area V5/MT remembers "what" but not "where". Cereb.Cortex 16(12), 1766–1770.

Campana G, Pavan A, Maniglia M, Casco C. (2011). The fastest (and simplest), the earliest: the locus of processing of rapid forms of motion aftereffect. Neuropsychologia. (10): 2929–34

Campana G, Maniglia M, Pavan A. (2013). Common (and multiple) neural substrates for static and dynamic motion after-effects: a rTMS investigation. Cortex. 49(9): 2590–4

Cattaneo, Z., & Silvanto, J. (2008). Investigating visual motion perception using the transcranial magnetic stimulation-adaptation paradigm. Neuroreport, 19(14), 1423–1427.

Cattaneo, Z., Silvanto, J., Battelli, L., & Pascual-Leone, A. (2009). The mental number line modulates visual cortical excitability. Neuroscience Letters, 462(3), 253–256.

Cattaneo, Z., Rota, F., Vecchi, T., & Silvanto, J. (2008). Using state-dependency of transcranial magnetic stimulation (TMS) to investigate letter selectivity in the left posterior parietal cortex: A comparison of TMS-priming and TMS-adaptation paradigms. European Journal of neuroscience, 28(9), 1924–1929.

Cattaneo, L. (2010) Tuning of ventral premotor cortex neurons to distinct observed grasp types: a TMS-priming study. Exp Brain Res., 207, 165–172.

Cohen-Kadosh, R., Muggleton, N., Silvanto, J, & Walsh, V. (2010). Double dissociation of format-dependent and number-specific neurons in human parietal cortex. Cerebral Cortex, 20(9), 2166–71.

de Graaf, T.A., Koivisto, M., Jacobs, C., & Sack, A.T. (2014). The chronometry of visual perception: review of occipital TMS masking studies. Neuroscience and Biobehavioral Reviews, 45, 295–304.

Guzman-Lopez, J., Silvanto, J., & Seemungal, B. M. (2011). Visual motion adaptation increases the susceptibility of area V5/MT to phosphene induction by transcranial magnetic stimulation. Clinical Neurophysiology, 122(10), 1951–1955.

Jacquet, P. O., & Avenanti, A. (2015). Perturbing the action observation network during perception and categorization of actions’ goals and grips: state-dependency and virtual lesion TMS effects. Cerebral Cortex, 25(3), 598–608.

Kammer, T. (2007). Visual masking by transcranial magnetic stimulation in the first 80 milliseconds. Advances in Cognitive Psychology, 3(1-2), 177–179.

Kammer, T., Puls, K., Erb, M., & Grodd, W. (2005). Transcranial magnetic stimulation in the visual system. II. Characterization of induced phosphenes and scotomas. Experimental Brain Research, 160(1), 129–140.

Kammer, T. (2007). Masking visual stimuli by transcranial magnetic stimulation. Psychological Research, 71(6), 659–666.

Kohn, A. (2007). Visual adaptation: physiology, mechanisms, and functional benefits. Journal of Neurophysiology, 97(5):3155–64.

Kohn, A., & Movshon, J. A. (2003). Neuronal adaptation to visual motion in area MT of the macaque. Neuron, 39(4), 681–691.

Mazzoni N, Jacobs C., Venuti P, Silvanto J, Cattaneo L. (2017). State-dependent TMS reveals representation of affective body movements in the anterior intraparietal cortex. J Neurosci. pii: 0913-17. doi: 10.1523/JNEUROSCI. 0913–17. 2017.

Moliadze, V., Zhao, Y., Eysel, U., & Funke, K. (2003). Effect of transcranial magnetic stimulation on single-unit activity in the cat primary visual cortex. The Journal of Physiology, 553(2), 665–679.

Nichols, M. J., & Newsome, W. T. (2002). Middle temporal visual area microstimulation influences veridical judgments of motion direction. The Journal of Neuroscience, 22(21), 9530–9540.

Renzi, C., Vecchi, T., Silvanto, J., & Cattaneo, Z. (2011). Overlapping representations of numerical magnitude and motion direction in the posterior parietal cortex: a TMS-adaptation study. Neuroscience Letters, 490(2):145–9.

Romei, V., Thut, G., & Silvanto, J. (2016) Information-Based Approaches of Noninvasive Transcranial Brain Stimulation. Trends Neurosci., 39, 782–795.

Rust, N. C, Mante, V., Simoncelli, E. P., & Movshon, J. A. (2006). How MT cells analyze the motion of visual patterns. Nature neuroscience, 9(11), 1421–1431.

Ruzzoli, M., Abrahamyan, A., Clifford, C. W., Marzi, C. A., Miniussi, C., & Harris, J. A. (2011). The effect of TMS on visual motion sensitivity: an increase in neural noise or a decrease in signal strength? Journal of Neurophysiology, 106(1), 138–143.

Schwarzkopf, D. S., Silvanto, J., & Rees, G. (2011). Stochastic resonance effects reveal the neural mechanisms of transcranial magnetic stimulation. The Journal of Neuroscience, 31(9), 3143–3147.

Siebner HR, Lang N, Rizzo V, Nitsche MA, Paulus W, Lemon RN, et al.. (2004). Preconditioning of low-frequency repetitive transcranial magnetic stimulation with transcranial direct current stimulation: evidence for homeostatic plasticity in the human motor cortex. J Neurosci. 24, 3379–85

Silvanto, J., Muggleton, N. G., Cowey, A., & Walsh, V. (2007). Neural adaptation reveals state-dependent effects of transcranial magnetic stimulation. European Journal of Neuroscience, 25(6), 1874–1881.

Stewart L, Battelli L, Walsh V, Cowey A (1999) Motion perception and perceptual learning studied by magnetic stimulation. Electroencephalogr Clin Neurophysiol Suppl 51:334–350

Solomon, S.G., & Kohn, A (2014) Moving sensory adaptation beyond suppressive effects in single neurons. Current Biology, 24 (20), 1012–1022.

Stocks, N. G. (2000). Suprathreshold stochastic resonance in multilevel threshold systems. Physical Review Letters, 84(11), 2310.

Stokes, M.G. (2015). "Activity-silent" working memory in prefrontal cortex: a dynamic coding framework. Trends in Cognitive Sciences, 19(7), 394–405.

Théoret, H., Kobayashi, M., Valero-Cabré, A., & Pascual-Leone, A. (2003). Exploring paradoxical functional facilitation with TMS. Supplements to Clinical Neurophysiology. 56, 211–219.

Théoret H, Kobayashi M, Ganis G, Di Capua P, Pascual-Leone A (2002). Repetitive transcranial magnetic stimulation of human area MT/V5 disrupts perception and storage of the motion aftereffect. Neuropsychologia.; 40(13):2280–7.

Walsh, V., & Pascual-Leone, A. (2003). Transcranial magnetic stimulation: a neurochromometrics of mind. Cambridge: MIT Press.

